# Unconscious integration of multisensory bodily inputs in the peripersonal space shapes bodily self-consciousness

**DOI:** 10.1101/048108

**Authors:** Roy Salomon, Jean-Paul Noel, Marta Łukowska, Nathan Faivre, Thomas Metzinger, Andrea Serino, Olaf Blanke

## Abstract

Recent studies have highlighted the role of multisensory integration as a key mechanism of self-consciousness. In particular, integration of bodily signals within the peripersonal space (PPS) underlies the experience of the self in a body we own (self-identification) and that is experienced as occupying a specific location in space (self-location), two main components of bodily self-consciousness (BSC). Experiments investigating the effects of multisensory integration on BSC have typically employed supra-threshold sensory stimuli, neglecting the role of unconscious sensory signals in BSC, as tested in other consciousness research. Here, we used psychophysical techniques to test whether multisensory integration of bodily stimuli underlying BSC may also occur for multisensory inputs presented below the threshold of conscious perception. Our results indicate that visual stimuli rendered invisible (through continuous flash suppression) boost processing of tactile stimuli on the body (Exp. 1), and enhance the perception of near-threshold tactile stimuli (Exp. 2), only once they entered peripersonal space. We then employed unconscious multisensory mechanisms to manipulate BSC. Participants were presented with tactile stimulation on their body and with visual stimuli on a virtual body, seen at a distance, which were either visible or rendered invisible. We report that if visuo-tactile stimulation was synchronized, participants self-identified with the virtual body (Exp. 3), and shifted their self-location toward the virtual body (Exp.4), even if visual stimuli were fully invisible. Our results indicate that multisensory inputs, even outside of awareness, are integrated and affect the phenomenological content of self-consciousness, grounding BSC firmly in the field of psychophysical consciousness studies.

## 1. Introduction

Based on clinical and experimental research in humans, it has been proposed that multisensory integration is a key mechanism for self-consciousness. In particular, bodily self-consciousness (BSC) has been shown to depend on the integration of multisensory bodily stimuli (Blanke, 2012; Blanke, Slater, & Serino, 2015; Ehrsson, 2012a; Tsakiris, 2010). Research has focused on two central aspects of BSC: people normally self-identify with a given body, which they perceive as their own (self-identification) and they experience their self at the location of their body (self-location) (Blanke, 2012; Blanke & Metzinger, 2009). Important support that BSC depends on multisensory integration of bodily inputs comes from research in neurological patients who suffer from alterations in the integration of such inputs leading to altered own body perceptions (Blanke, Landis, Spinelli, & Seeck, 2004; Blanke, Ortigue, Landis, & Seeck, 2002). Another key demonstration was provided by experimental manipulations of BSC in healthy subjects using multisensory conflicts (Ionta et al., 2011; Lenggenhager, Tadi, Metzinger, & Blanke, 2007; Petkova & Ehrsson, 2008; Petkova, Khoshnevis, & Ehrsson, 2011; Salomon, Lim, Pfeiffer, Gassert, & Blanke 2013). For example, in the full body illusion, viewing an avatar’s body being stroked, while concurrently receiving the same tactile stimulation on one’s own body, makes participants self-identify with the avatar (Ehrsson, 2007; Petkova & Ehrsson, 2008) and induces changes in self-location such that subjects perceive themselves closer to the avatar’s position (Ionta et al., 2011; Lenggenhager et al., 2007).

Under normal conditions, multisensory body-related stimuli occur within a limited distance from the body, which defines the peripersonal space (PPS). Accordingly, neuronal populations have been described both in monkeys and in humans integrating somatosensory stimulation on the body with visual and/or auditory stimuli specifically when presented close to the body (Graziano & Cooke, 2006; Ladavas & Serino, 2008; Rizzolatti, Fadiga, Fogassi, & Gallese, 1997). PPS and BSC share common neural structures in premotor, posterior parietal, and temporo-parietal cortex (Blanke et al., 2015; Makin, Holmes, & Ehrsson, 2008) and it has recently been shown that the full body illusion leads to a shift in PPS from the physical body toward the virtual body that participants identify with, compatible with an extension of the PPS boundary (Noel, Pfeiffer, Blanke, & Serino, 2015; Serino, Canzoneri, Marzolla, di Pellegrino, & Magosso, 2015). These data link processing and integration of multisensory stimuli within PPS to self-consciousness, and to BSC in particular (Blanke et al., 2015).

Conscious experience has also been related to the integration of sensory information in the brain by other authors (Dehaene & Naccache, 2001; Mudrik, Faivre, & Koch, 2014; Tononi, 2008). Indeed, consciousness is characterized by a unity of experience in which information from multiple sensory modalities is integrated and bound together (Bayne, 2002; James, Burkhardt, Bowers, & Skrupskelis, 1981). Recent experimental work has shown that consciously perceived non-visual stimuli may even be integrated with stimuli rendered invisible through various masking paradigms (i.e. auditory (Alsius & Munhall, 2013; Lunghi, Morrone, & Alais, 2014), tactile (Lunghi & Alais, 2013; Lunghi, Binda, & Morrone, 2010; Salomon, Galli, et al., 2015), olfactory (Zhou, Jiang, He, & Chen, 2010), proprioceptive (Salomon, Lim, Herbelin, Hesselmann, & Blanke, 2013) and vestibular (Salomon, Kaliuzhna, Herbelin, & Blanke, 2015)) and that even a subliminal auditory and a subliminal visual stimulus can be integrated and impact consciousness (Faivre, Mudrik, Schwartz, & Koch, 2014; Noel, Wallace, & Blake, 2015). Do these findings on unconscious integration also extend to self-consciousness and BSC in particular, which is often considered a more complex and specific form of conscious content (Dehaene & Changeux, 2011; Faivre, Salomon, & Blanke, 2015; Gallagher, 2000)?

Previous research on the multisensory basis of BSC focused on the integration of sensory inputs that are presented above participants’ visual and tactile thresholds. Yet as it has been argued that BSC is based on low-level and pre-reflexive brain mechanisms, it is possible that the sensory events shaping the experience of the self need not be consciously perceived. However, to date there is no experimental evidence suggesting that the multisensory integration processes of BSC do not require conscious awareness of the respective multisensory stimuli, although unconscious multisensory integration has been shown in humans (see above) (Faivre et al., 2014; Salomon, Kaliuzhna, et al., 2015; Salomon, Lim, Herbelin, et al., 2013) and at the neuronal level in anesthetized animals (Graziano, Hu, & Gross, 1997; Meredith & Stein, 1986; Stein & Stanford, 2008). Here, we tested for the first time whether multisensory integration of bodily stimuli underlying BSC also occurs for multisensory inputs, which are presented below the threshold of conscious perception. To this aim, in a series of four experiments, we first tested the hypothesis that multisensory integration of body-related signals within the PPS occurs also for stimuli presented below the threshold of conscious perception. We asked whether tactile stimuli on the body are preferentially integrated with visual stimuli presented within, as compared to outside the PPS, when visual inputs were subliminal and tactile inputs supraliminal (Exp. 1) or even when visual was subliminal and tactile inputs were near-threshold (Exp. 2). Next, we investigated whether it is possible to manipulate BSC by using visuo-tactile conflicts administered below the perceptual threshold. To this aim, we coupled tactile stimulation on the body with invisible synchronous visual stimuli on a virtual body to induce the full body illusion (Lenggenhager et al., 2007) and tested whether this would affect self-identification, as assessed by questionnaires (Exp.3) and self-location, as assessed by the location of PPS boundaries (Exp. 4).

## 2. Methods

### 2.1 Participants

In total 98 participants (31 females, mean age = 23.0 ± 2.7) were included in this series of experiments. Sample sizes were determined based on the effect sizes of our prior work (Noel et al., 2015; Serino et al., 2015). Thirty-two subjects took part in Exp. 1, 15 in Exp. 2, 25 in Experiment 3, and 26 in Exp. 4 (the first experiment being a between-subject experimental design, while the latter three being within-subjects). All participants were right-handed, had normal or corrected-to-normal visual acuity, reported normal hearing and touch, and had no history of psychiatric or neurological disorder. All volunteers provided written informed consent to participate in the study, which was approved by the Brain Mind Institute Ethics Committee for Human Behavioral Research of the EPFL, and conducted in accordance with the Declaration of Helsinki.

### 2.2 Materials and Procedure

#### 2.2.1 Experiment 1

Visual stimuli consisted of a three-dimensional virtual white wireframe ball either looming toward or receding from the participants’ face (Fig 1A). The ball, presented in stereoscopy, travelled approximately 2 meters in virtual space at a velocity of 50 cm/s until making fictive contact with the participant’s face, or in the opposite direction in the case of receding stimuli.

**Fig. 1.**
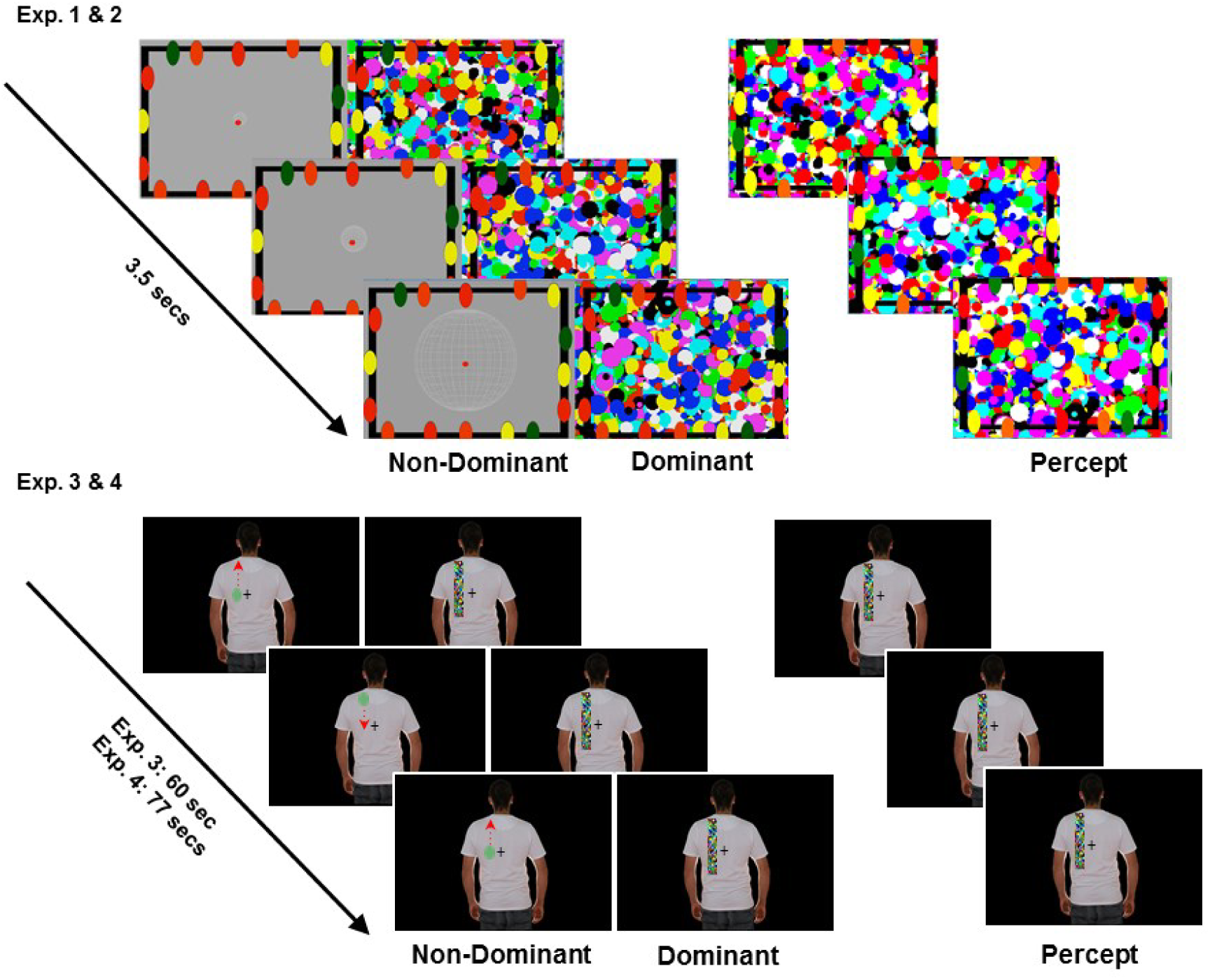
Experimental design. Top. Experimental stimuli in the Invisible condition in Exp. 1 & 2. A wireframe ball approaching the participants’ face was presented to the non-dominant eye while highly salient colored masks were rapidly (10hz) flashed to the dominant eye (CFS masking). Due to CFS, participants perceived the masks, while the approaching ball was invisible. Bottom. Experimental stimuli in the Invisible condition in Exp. 3 & 4. An image of a body with a moving dot on the back was presented to the non-dominant eye. The dot could be moving synchronously or asynchronously to the tactile stimulation on the participants’ back. Critically, CFS masking of the region of the dot movement in the Invisible trials rendered the dot invisible, thus, both in the synchronous and asynchronous stimulation condition the percept was of a body image with rectangular flashing masks only.

Visual stimuli were presented on a head-mounted display (HMD, VR1280 Virtual Research Systems, Inc., Santa Clara, CA, USA) with a resolution of 1280x1024 pixels, representing a 60-degree diagonal field of view, at 60 Hz. Participants in the Invisible group, were also presented with circular high-contrast dynamic noise patches suppressors (“Mondrians”;(Salomon, Galli, et al., 2015)). These suppressors were rapidly (10 Hz) flashed to the participants’ dominant eye (determined by the experimenter via the Miles test (Miles, 1930)prior to the study).

In addition to the visual stimuli, participants’ were outfitted with a vibrotactile device (Precision MicroDrives shaftless vibration motors,), placed on the subjects’ forehead. Vibrotactile stimulation was presented supra-threshold for a duration of 100 milliseconds. Participants provided responses to vibrotactile stimulation with a wireless gamepad (XBOX 360 controller, Microsoft), which they held in their right. In-house software ExpyVR (freely available at http://lnco.epfl.ch/expyvr) was utilized for the rendering and presentation of visual and vibrotactile stimuli. RTs were measured relative to the onset of tactile stimulation.

On experimental trials (70% of total trials), tactile stimulation was administered above-threshold via a vibrotactile device on the face. Concurrently to the tactile stimulation, a looming or receding visual stimulus was presented via a Head-Mounted Display (HMD). Half the participants performed the task while the visual stimuli presented was visible, whereas for the other half of participant the dynamic visual stimulus was suppressed via Continuous Flash Suppression (CFS; Tsuchiya & Koch, 2005). Visuo-tactile interactions were probed through space at 7 different visuo-tactile distances (D1 through D7). That is, after initiation of a trial, dynamic visual stimuli were shown as either approaching or receding from the subject, and, at a particular temporal delay (T1 through T7), tactile stimulation, to which participants were to respond, was administered (see Supplementary Information for details). Baseline trials (20% of total trials) – trials in which no visual stimulation was given – were also sampled at T1 and T7 in order to measure unimodal tactile RT; these data were used to correct for a spurious temporal effect and in order to confirm that speeding in RTs as a consequence of visual stimuli within PPS reflected true multisensory facilitation.

#### 2.2.2 Experiment 2

Materials and procedure followed as for Experiment 1, with two exceptions. First, visible and invisible conditions of visual stimulation were administered within-subjects, in separate blocks, whose administration order was counter-balanced between participants. Second, tactile target stimuli were presented at perceptual threshold by means of masking tactile stimuli. The tactile target stimulus was given by means of a miniature solenoid, whereas the masking administered by means of 4 vibrotactile stimulators, placed surrounding the solenoid and activated throughout the duration of a trial. The intensity of the tactile target stimulus on the face was titrated (by means of a staircase procedure) before each experimental block so to be detected on 60% of trials, without visual stimulation (see Supplementary Information for further details).

#### 2.2.3 Experiment 3

The procedure to induce the full body illusion consisted in applying tactile stimulation on the participants back and visual stimulation on a virtual body (avatar; H: 20,5° W: 11,3°), seen through a HMD. Tactile stimulation was administered by using a haptic robotic system (Salomon, Lim, Pfeiffer, et al., 2013). Visual stimuli (see Fig. 1B) consisted of a colored visual dot (size: H: 0.7°, W: 0.7) that was moving up and down along the left side of the avatar’s back. In the critical condition inducing the illusion, the movement of the haptic robot was fully synchronized temporally and spatially with that of the dot on the avatar’s back. In the control, asynchronous condition, the visual and tactile stimulation were uncorrelated by using unmatched visual and tactile motion profiles. In order to make the pattern of visuo-tactile stimulation invisible to the participants, visual stimuli was administered in a CFS paradigm, whereby the visual dot was presented to the non-dominant eye, while to the dominant eye a stream of high-contrast dynamic noise patch suppressors (“Mondrians”, as described in(Salomon, Galli, et al., 2015);(H:8,9°, W: 1°)) was presented, on a rectangular section on the left of the avatar’s back with a frequency of 10Hz (see Fig.1B). After each 60 seconds of visuo-tactile stimulation, participants responded to questions regarding masking efficiency (see Supplementary Information). In order to measure the effect of the illusion at the subjective level, two testing self-identification and referred touch were administered.

#### 2.2.4 Experiment 4

The procedure to induce the full body illusion was identical to that of experiment 3, with two differences: the omission of the non-masked (visible) condition and longer visuo-tactile stimulation lasting 77 seconds per trial, allowing intermingled testing of PPS. In order to assure that participants were not aware of the pattern of visuo-tactile stimulation, participants were required on each trial to press a button in case they saw the dot. Intermingled with visuo-tactile stimulation, PPS was measured via an audio-tactile paradigm (21, 48, 49). The task was similar to that described for Experiment 1 and 2, with the exception that an auditory (broadband noise), and not a visual stimulus approached the participant’s chest. Six different audio-tactile distances were probed (see Supplementary Information online). We used audio-tactile stimulation, instead of visuo-tactile stimulation (as in Exp. 1 & 2), in order to keep the experimental manipulation used to induce the full body illusion (visuo-tactile stroking) and that used to measure its effect on peripersonal space (audio-tactile interaction) orthogonal with each other (as in Noel et al., 2015).

### 2.3 Data Analyses

Trials in which participants reported seeing the visual stimuli, correctly identified the color or did not respond to the awareness questions were removed from the analysis (28% of trials in Exp. 1 and 21% in Exp.2; 4% in Exp. 3 and 12% in Exp. 4).

For PPS measurement (Exp. 1-2 & 4), we first calculated on a subject-per-subject basis the mean RT (Exp. 1 & 4) and detection rates (Exp. 2) for the baseline unimodal tactile conditions. Subsequently, the fastest mean baseline condition (i.e., T1) was subtracted from the participant mean in all the other conditions to provide a measure of facilitation induced on tactile processing by visual or auditory stimuli perceived at a different distance from the participant’s body (See (Noel et al., 2014; Noel et al., 2015) for a similar approach). Subsequently, on a subject-per-subject basis, RT or detection rates relative to baseline were fitted to both linear and sigmoidal curves (see Canzoneri, Magosso, & Serino, 2012, for details). For each experiment we modeled the data with the best fit (linear for Exp. 1 & 2, and sigmoidal in Exp. 3 – See Supplementary Information) and then compared across conditions the values extracted from the fitting procedure. In experiment 3, we analyzed responses to BSC questions (Q1 & Q2) during the visible and invisible conditions and using a repeated measures ANOVA with synchronicity (Synchronous/Asynchronous) and visibility (Visible /Invisible) as within-subject factors. When interactions were present, t-tests were used to explore modulation of BSC within each synchronicity level.

## 3. Results

### 3.1 Invisible looming stimuli within the PPS affect tactile perception (Exp 1)

In Experiment 1, participants were asked to respond as fast as possible to an above threshold tactile stimulation administered on their face. On experimental trials (70%), they concurrently received a task-irrelevant visual stimulus, administered through a head-mounted display (HMD), consisting of a virtual ball approaching their face. In two conditions, the visual stimulus was either clearly perceived (visible condition), or was rendered invisible (invisible condition) by using continuous flash suppression, a well-established psychophysical technique in which highly salient mask images (“Mondrians”) presented to the dominant eye suppress awareness of a target image presented to the other eye for an extended period of time (Tsuchiya & Koch, 2005; Yang & Blake, 2012). In the invisible condition, the virtual ball was presented to the non-dominant eye, while Mondrians were concurrently flashed to the dominant eye, whereas in the visible condition, the virtual ball was presented to both eyes (Fig. 1A: see Supplementary Information online for a full description of the continuous flash suppression procedure and control experiments). We delivered the tactile stimulus, to which participants were asked to respond, at seven different time delays from the onset of visual stimulation. In this way tactile stimuli were associated with visual stimuli presented at 7 different distances from the participant’s body (from close (D1) to far (D7)) allowing us to test whether tactile detection is tuned to visual distance. Previous studies using the same protocol showed that reaction times (RT) to tactile stimulation decrease once a stimulus enters the participant’s PPS (Canzoneri et al., 2012; Teneggi, Canzoneri, di Pellegrino, & Serino, 2013). Here we investigated whether the distance-dependent modulation of tactile RT is present even when the approaching visual stimuli entering PPS are invisible, suggesting that multisensory integration within the PPS occurs also in the absence of visual awareness. In order to control for a mere temporal effect (i.e., participants might become faster at later delays), we also included a control condition, whereby receding visual stimuli were administered, and for which we predicted no distance-dependent modulation of RT for face stimulation (Serino et al., 2015; Teneggi et al., 2013).

We analyzed RT to the tactile stimulation as a function of the different distances of the virtual ball and its direction, in the visible and invisible conditions. As shown in Fig. 2A, there was a clear distance dependent modulation of RT, as a function of the location of the visual stimulus, both for the Visible and Invisible conditions. This was not the case for Receding visual stimuli, excluding that the present finding was a mere temporal effect (see supplementary material online, Fig. S2A). Next, we fitted individual data to a linear function (which was the model to best fit the results; see supplementary analysis online), comparing the slope of the function, as a measure of how strongly tactile processing was influenced by the location of the task-irrelevant visible and invisible approaching balls. The presence of a positive slope, steeper for looming visual stimuli, would indicate a stronger multisensory integration effect for visual stimuli entering the PPS. The slope values were submitted to a 2X2 mixed ANOVA with Ball Direction (Looming and Receding), as within-subjects factor, and Condition (Visible and Invisible), as between-subject factor. The main effect of Ball Direction was significant (F(1,28)=69.52, p<.0001, partial η^2^=0.71): the slope of the function was positive only for looming (mean slope=0.33±.02) and not for receding (mean slope=0.07±.02) stimuli. There was no main effect of Condition (p=0.64,1–β= 0.7), nor a Condition X Ball Direction interaction (p=0.93,1–β=0.5). Thus, the modulation of tactile processing due to the distance of the task-irrelevant visual stimuli at the time of touch was found for both visible and invisible balls. Importantly, the positive value of the looming slope was significantly different from zero for both conditions (visible: t(14)=11.80,p<0.001; invisible: t(16)=11.60,p<0.001). Thus, a distance-dependent modulation of tactile processing was found when task-irrelevant looming stimuli, that were not consciously perceived, were presented, indicating that multisensory integration within the PPS occurs even in absence of awareness for the visual stimulus.

**Fig. 2.**
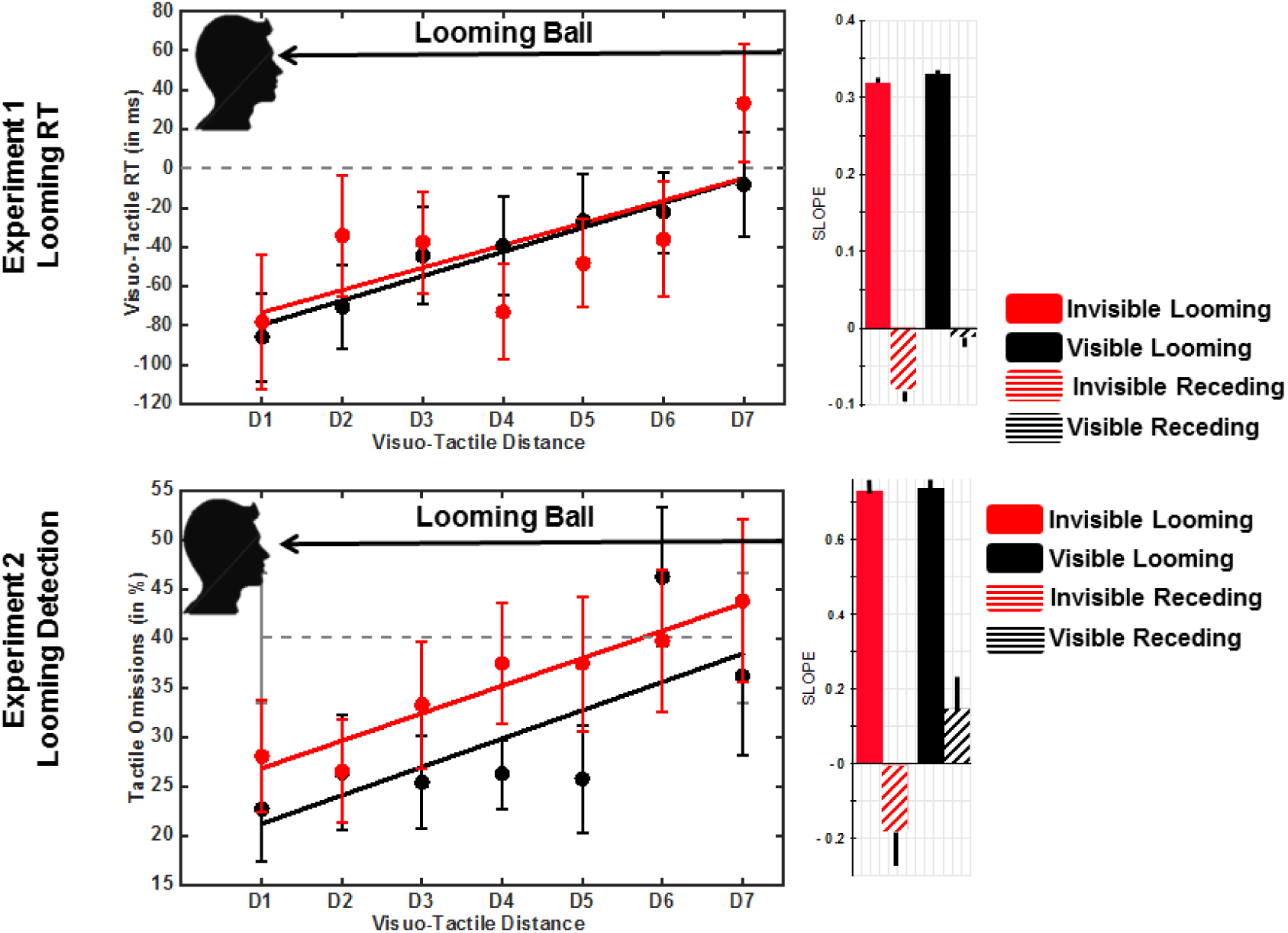
PPS in absence of awareness. A) Experiment 1. RTs to tactile targets as a function of the distance of the approaching visual stimulus. In order to show a truly multisensory visuo-tactile facilitation effect, RTs are reported as the difference between responses to tactile stimuli when they were coupled with visual stimulation and responses to tactile stimulation alone. Baseline unimodal tactile RTs (administered in 20% of trials) are thus by definition equal to zero (illustrated by the dashed line; (J. P. Noel et al., 2015)). Data for both the conditions in which the looming visual stimuli was visible (black) and invisible (red) were fitted to linear functions (see supplementary material online). Error bars indicate +/- 1 S.E.M. For both conditions, tactile processing speeded up as the visual stimulus approached the body. B) Exp. 2. Omission to tactile targets as a function of the distance of the approaching visual stimulus (Convention follows as in A). Tactile stimulation was set to be detected on 60% of trials, (i.e., omitted on 40% of unimodal tactile trials). Perception increased as the ball approached the body, both in the visible (black) and invisible (red) conditions.

### 3.2 Invisible looming stimuli increase tactile awareness (Exp 2)

In Experiment 2, we investigated whether invisible visual stimuli, occurring within the PPS, modulate not only the processing of supra-threshold tactile stimuli, but also enhance the perception of near-threshold tactile stimulation. To this aim, we used a staircase procedure (see Supplementary Information online), so that tactile targets were perceived in 60% of trials, when presented alone. Then, near-threshold tactile target stimuli were coupled with looming (or receding, as a control condition) visual stimuli that were again either fully visible or rendered invisible through CFS (as in Exp. 1). We predicted that visible and invisible visual stimuli occurring within PPS would also boost the detection of near-threshold tactile stimuli (but only for looming stimuli), thus increasing subjects’ accuracy in reporting tactile stimulation. Fig.2B reports the percentage of missed tactile targets as a function of the distance of looming visual stimuli and shows that tactile detection increased as the virtual ball approached the subjects (see Fig.2B). Data were fitted with a linear function (as the best model fitting the data, see supplementary analyses online) and analyzed as in Exp. 1. The main effect of Ball Direction was significant (F(1,14)=287.03,p< 0.001,partial η^2^=0.95), with steeper slopes for looming (mean slope=0.73±0.03) as compared to receding visual stimuli (mean slope=0.07±0.001) (see Fig. S2B). As in Exp. 1, there was no main effect of Condition (p= 0.31,1 –β=0.66), nor a Condition X Ball Direction interaction (p=0.18,1–β=0.55), meaning that the same spatially dependent modulation of tactile perception was found both in the visible and in the invisible conditions. To summarize, visual stimuli within the PPS, enhance the perception of near-threshold tactile stimuli on the body, even when they are rendered fully invisible.

### 3.3 Invisible visuo-tactile conflicts modulate self-identification (Exp 3)

Having demonstrated visuo-tactile integration for unconscious sensory inputs within PPS, we next asked whether we could modulate BSC by manipulating the spatio-temporal congruency of visuo-tactile stimuli (Blanke, 2012; Ehrsson, 2007; Lenggenhager et al., 2007), even when the multisensory conflict was not consciously perceived. To this aim, in Exp. 3, we used visuo-tactile stimulation to induce the full body illusion using either fully visible stimuli (as done in previous studies) or identical visual stimuli rendered invisible by means of CFS. Participants received above-threshold tactile stimulation on their back, administered by means of a robotic stroking set up (Ionta et al., 2011; Salomon, Lim, Pfeiffer, et al., 2013), while concurrently seeing an avatar from behind, presented binocularly through a head-mounted display. The avatar was shown on the HMD as receiving tactile stimulation on the back, represented by a colored dot moving at the same speed and to the same extent as the tactile stimulation participants received on their back (see Fig. 1B). In the synchronous condition, normally inducing the full body illusion (Ionta et al., 2011; Lenggenhager et al., 2007; Salomon, Lim, Pfeiffer, et al., 2013), the visual stimulation on the avatar’s body and tactile stimulation on the participant’s body were corresponding. An asynchronous visuo-tactile stimulation, in which the visual and tactile stimulations were unrelated, was administered as a control condition. The experiment was run in a 2X2 factorial design, whereby beyond synchrony of stimulation, we also manipulated visibility of the moving dot, which was either fully visible, as in the standard full body illusion, or rendered invisible by masking the region of visual stroking with Mondrian patterns flashed to the dominant eye (see Supplementary Information online and (Salomon, Galli, et al., 2015) for details). On each trial, participants were stroked for one minute. Stimulus visibility was vigorously controlled (see Supplementary Information for full details). Trials in which participants reported seeing a visual stimulus apart from the masks were removed from analysis (3% of trials). In the remaining fully suppressed trials participants were at chance for reporting the dot’s color and visuo-tactile synchrony (mean accuracy 49% and 50% respectively see supplementary materials for further analysis). The modulation of BSC was measured with two questions (modified from (Lenggenhager et al., 2007)) probing self-identification (Q1: *‘How strong was the feeling that the body you saw was you?’*) and illusory touch (Q2: *‘How strong was the feeling that the touch you felt originated from the body you saw?’*), using a scale from 1 (*Completely disagree*) to 10 (*Completely agree*).

Participants’ responses indicated that a change in BSC was obtained by means of synchronous stimulation both in the visible and in the invisible conditions. First, a repeated measures ANOVA on Q1 scores with synchrony (Synchronous/Asynchronous) and visibility (Visible/Invisible) as within-subject factors revealed a significant main effect of synchrony (F(1,19)=24.47,p=.00009, partial η^2^=0.56), with higher self-identification in the synchronous (M=4.0, S.E.M=0.59) than in the asynchronous (M=3.2, S.E.M=0.59) condition. Moreover, the main effect of visibility was significant (F(1,19)=8.08, p=.01, partial η^2^=0.29), with higher self-identification ratings in the visible (M=4.1, S.E.M=0.55) than in the Invisible (M=3.0, S.E.M=0.45) condition. The interaction between synchrony and visibility was also significant (F(1,19)=7.41, p=0.014, partial η^2^=0.28), with larger differences in self-identification as a function of synchrony ratings in the visible (Visible-synchronous M=4.8, S.E.M=0.56, Visible-asynchronous M=3.5, S.E.M=0.51) than the invisible (Invisible-synchronous M=3.2, S.E.M=0.44, Invisible-asynchronous M=2.8, S.E.M=0.48) condition. Importantly, paired samples t-test revealed significantly higher ratings for self-identification with the avatar after synchronous as compared to asynchronous visuo-tactile stroking both in the Invisible (t(19)=2.31; p=0.03 two-tailed, Cohen’s *d*=0.54) and the Visible (t(19)=4.31; p=0.0001,Cohen’s d=1.02) (see Fig. 3A) condition. This result shows that visuo-tactile stimulation led to higher explicit self-identification responses in a synchrony-dependent manner even when participants were not aware of the type of visual stimulation they were receiving. Responses to the second question regarding illusory touch, revealed a significant main effect of synchrony, with higher misattribution of touch (F(1,19)=23.89,p=0.0001,partial η^2^=0.55) in the synchronous (M=3.3, S.E.M=0.45) than in the asynchronous (M=2.5, S.E.M=0.41) condition. The main effect of Visibility was not significant (F(1,19)=0.2, p=0.87). The interaction between visibility and synchrony was significant (F(1,19)=12.23, p=0.002, partial η^2^=0.39), with larger differences in illusory touch as a function of synchrony in the visible (Visible-synchronous M=3.6, S.E.M=0.56, Visible-asynchronous M=2.3, S.E.M=0.45) than the invisible (Invisible-synchronous M=3.6, S.E.M=0.56, Invisible-asynchronous M=2.3, S.E.M=0.45) condition. Importantly, as for self-identification, paired samples t-test indicated that participants misattributed tactile stimulation to the virtual body significantly more strongly in the case of synchronous as compared to asynchronous stimulation not only in the visible (t(19)=4.61; p=0.00009, Cohen’s d=1.07), but even in the invisible (t(19)=2.14; p=0.02 one-tailed, Cohen’s d=0.47) condition, i.e. when they were not aware of the spatio-temporal pattern of visuo-tactile stimulation (see Fig. 3A). Together, these findings show that modulations of BSC by visuo-tactile conflict occur even when the visual stimuli, and the resulting multisensory conflict, are not consciously experienced. This result is the first empirical evidence that explicit changes in the phenomenal content of BSC arise by manipulating multisensory cues in the absence of awareness.

**Fig. 3.**
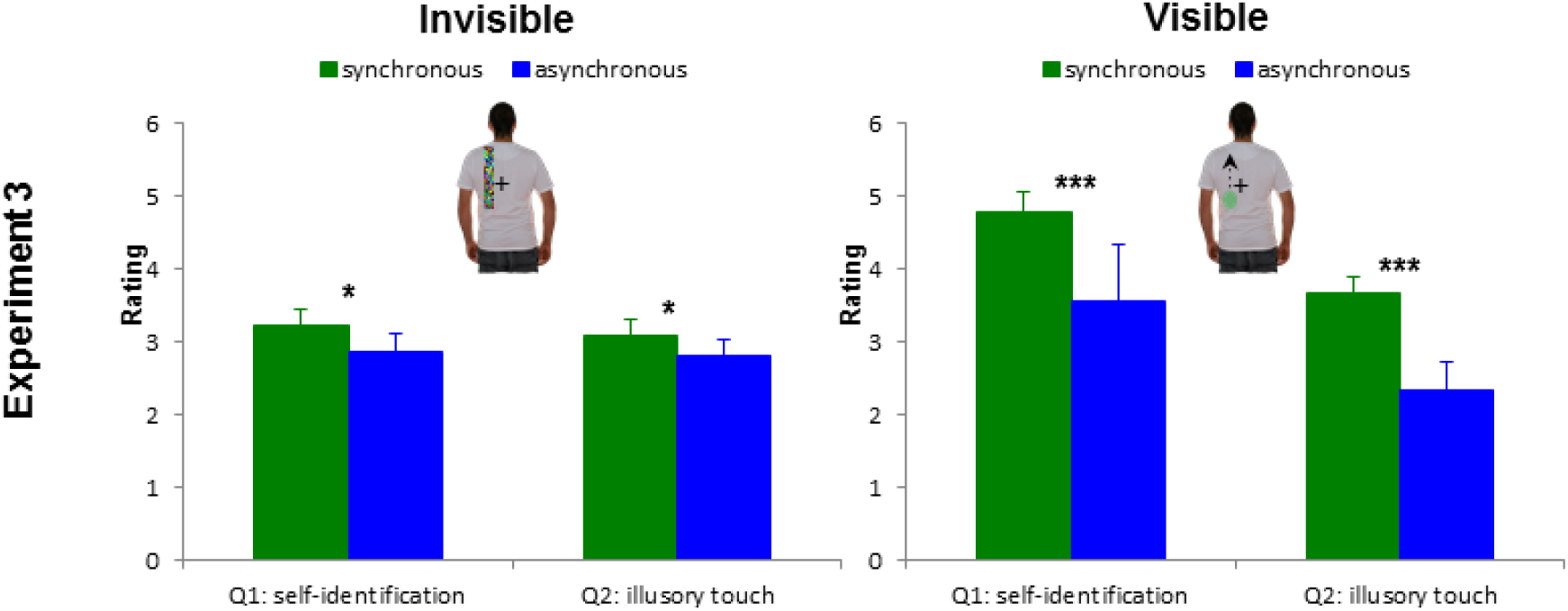
Modulation of self-identification by an invisible multisensory conflict. Responses to BSC questions relating to self-identification and illusory touch for synchronous and asynchronous visuo-tactile stimulation. Significant modulation was found for the full body illusion condition (synchronous visual tactile stimulation) for both invisible (left) and visible (right) conditions (within-subject error bars were calculated according to Cosineau method).

### 3.4 Invisible visuo-tactile conflicts modulate perceived self-location (Exp 4)

We finally investigated if an unconscious multisensory manipulation of BSC would also modulate self-location (Blanke, 2012; Lenggenhager, Mouthon, & Blanke, 2009; Lenggenhager et al., 2007). Previously, we showed that during the full body illusion (induced with fully perceived visual and tactile stroking), the boundaries of PPS representation, as assessed by means of an audio-tactile interaction task, shifted from being centered at the participants’ body, toward the location of the avatar’s body with whom the participants identified (Noel et al., 2015). Here, we applied the same paradigm, but tested whether a similar change in PPS, reflecting a change in self-location, can be achieved when visuo-tactile stimulation applied to induce the full body illusion is not visible to the participant. To this aim, epochs of masked visuo-tactile stimulation (as in Exp. 3) were intermingled with audio-tactile trials measuring PPS (see Methods and supplementary information for details). Perceptual awareness for the visual stimuli was controlled as in Exp. 3 and only trials in which the participants were completely unaware were included in the analysis (12% of trials were excluded, see Supplementary Information). The PPS paradigm was similar to that used in Exp1 of the present study, but we used auditory looming stimuli, instead of visual stimuli, in order to keep the form of multisensory stimulation used to induce the full body illusion (visuo-tactile) orthogonal to that used to test its effect on perceived self-location (auditory-tactile). Participants were requested to respond as quickly as possible to a tactile vibration administered on their trunk, while task-irrelevant sounds approached their body. Figure 4A shows RT to tactile targets as a function of the distance of the sound at the time of tactile stimulation. In order to test whether the boundaries of PPS varied between the synchronous and the asynchronous stroking conditions, RTs were fitted with a sigmoidal function (Canzoneri et al., 2012; Serino et al., 2015; Teneggi et al., 2013). The sigmoidal’s central point, representing an index of the location of PPS boundary, and slope, representing an index of the gradient of PPS representation were compared (Synchronous vs. Asynchronous). The central point location was significantly different in the Synchronous (M = 4.5, S.E.M. = 0.22) as compared to the Asynchronous (M = 3.8, S.E.M. = 0.45) condition (t(20) =2.452, p = 0.024, partial η^2^=0.198), indicating that participants’ PPS boundary was more distant from the participant’s body, and thus closer to the avatar’s body, in the Synchronous condition than in the Asynchronous control condition. No synchrony effect was found on the slope (p=0.34, 1 – β = 0.73), which was however different from 0 in both conditions (both p-value<0.03), indicating a distance-dependent modulation of tactile processing. Thus, the manipulation of multisensory cues, of which participants were not aware of (yet inducing changes in the phenomenal content BSC, Exp. 3), caused a shift in self-location toward the virtual body participants identified with, as shown here based on the effect on the PPS boundary (Noel et al., 2015).

**Fig. 4.**
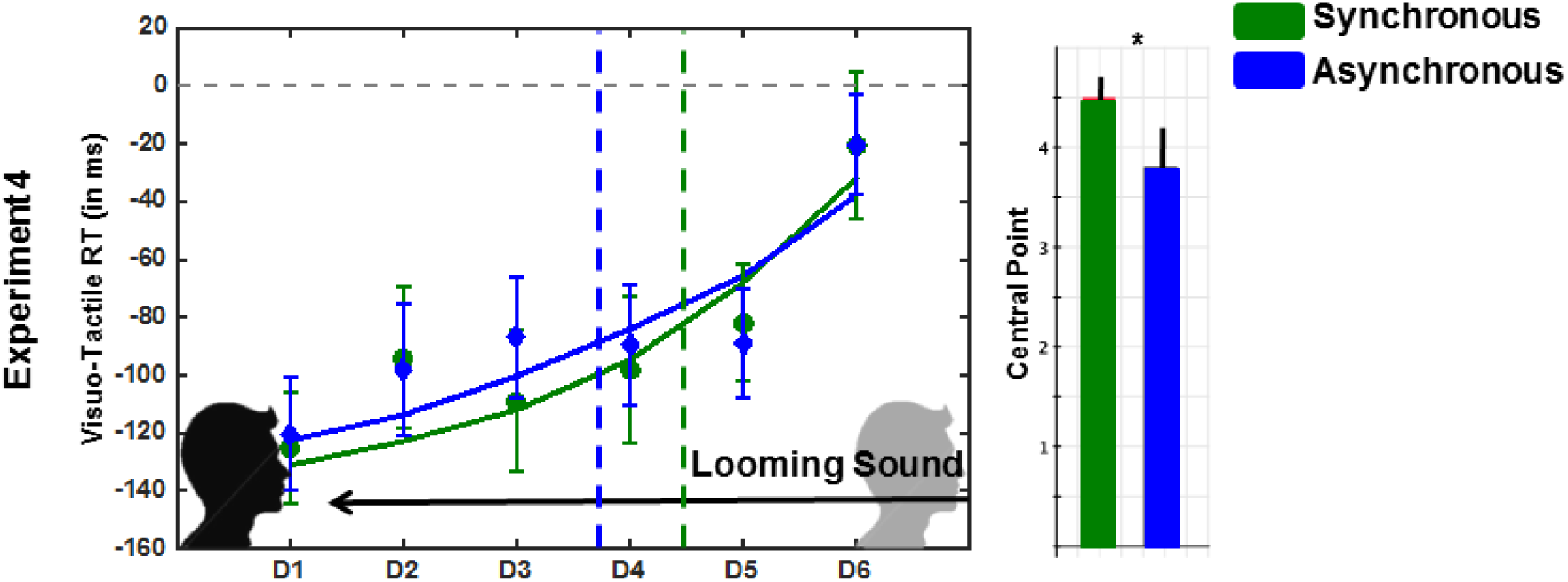
Modulation of self-location by an invisible multisensory conflict. RTs to tactile targets as a function of the distance of the approaching auditory stimuli (D7-D1) and the visuo-tactile stroking condition (synchronous in green and asynchronous in blue). RTs are reported as the difference between responses to tactile stimuli when they were coupled with visual stimulation and response to tactile stimulation alone. Baseline unimodal tactile RTs (administered on 20% of trials) are thus by definition equal to zero (illustrated by the dashed line). Data was fitted to a sigmoidal function. Error bars indicate +/- 1 S.E.M. The vertical dashed lines indicate the mean central point of the sigmoidal fitting, computed as a measure of the distance at which sounds start affecting RTs and analyzed in order to quantify PPS boundaries. This value was located at a farther distance in the synchronous (green) as compared to the asynchronous (red) visuo-tactile stroking conditions, indicating a more extended PPS in the former case.

## 4. Discussion

### 4.1 Unconscious multisensory integration in PPS

The self is essential to our understanding of consciousness (Blanke & Metzinger, 2009; Damasio, 2012; Metzinger, 2004) and recent work has highlighted the role of multisensory integration and PPS in self-consciousness, especially in BSC (for reviews see Blanke, 2012; Blanke et al., 2015; Ehrsson, 2012b; Noel et al., 2015). The present study brings novel comprehensive evidence that multisensory integration in PPS does not require conscious awareness and, importantly, that these unconscious multisensory processes modulate the phenomenological content of BSC.

In the first two experiments we show that multisensory integration of bodily signals within the PPS occurs when visual stimuli are presented below the perceptual threshold. This was demonstrated by showing that visuo-tactile interaction in PPS occurs when visual stimuli are rendered invisible (Exp.1 & 2) and even when the tactile stimuli (associated with invisible visual stimuli) were presented near the tactile threshold (Exp.2). Thus, conscious perception of visual and tactile stimuli is not required for multisensory integration of bodily signals within the PPS.

Previous behavioral findings showed that the processing of invisible stimuli is affected by concurrent non-visual stimuli above perceptual threshold (Alsius & Munhall, 2013; Lunghi et al., 2010; Lunghi et al., 2014; Maruya, Yang, & Blake, 2007; Salomon, Lim, Herbelin, et al., 2013; Zhou et al., 2010). Data from experiment 1 demonstrate the complementary effect, in which invisible visual stimuli impact processing of supra-threshold tactile stimuli.

Experiment 2 further extends this by showing that an invisible visual stimulus even modulates awareness for tactile stimuli near the tactile threshold, thus extending recent work revealing interactions between two unconscious stimuli (auditory-olfactory (Arzi et al., 2012); auditory-visual (Faivre et al., 2014)). The present study is the first report, to the best of our knowledge, of a multisensory interaction between near-threshold tactile and visual stimuli and in revealing that this unconscious visuo-tactile effect depends on the distance from the body (PPS), compatible with findings in neurophysiological studies showing PPS-dependent responses in bimodal and trimodal neurons in anesthetized monkeys (Graziano, Hu, & Gross, 1997; Stein & Stanford, 2008).

### 4.2 Unconscious multisensory integration underlies BSC

Recent accounts suggest that modulation of BSC through manipulation of multisensory inputs, as during the full body illusion, depends on the extension of the visual receptive fields of bimodal PPS neurons (Blanke, 2012; Ehrsson, 2012b; Makin et al., 2008; Noel et al., 2015). Based on this and the findings of experiments 1 and 2, we predicted that sub-threshold multisensory stimulation may also impact BSC and subjective responses about the self. However, previous studies using visuo-tactile stimulation to manipulate BSC applied stimuli well above the perceptual thresholds (e.g. Ehrsson, 2007; Lenggenhager et al., 2007; Petkova & Ehrsson, 2008; Salomon, Lim, Pfeiffer, et al., 2013). While it is evident that we are not consciously aware of most multisensory states (including those of BSC), to date it is not known whether unconscious multisensory stimuli can influence the content of BSC and how such effects with unconscious stimulation compare to effects obtained with conscious stimulation. Here we show that subjective and objective responses about the phenomenal content of BSC are modulated by unconscious multisensory stimulation and that this modulation, although weaker, is qualitatively comparable to modulations obtained with fully conscious stimuli. Experiment 3 indicated that for two patterns of stimulation, which were perceptually identical to the participants – i.e., seeing an avatar (without seeing the stroking) and feeling tactile stimulation, different explicit self-related experiences were induced that depended on an unperceived temporal relationship between visual and tactile stimulation (i.e., synchronous vs. asynchronous). In experiment 4 we show that this unconscious multisensory integration not only alters self-identification, but also impacts self-location, as we observed a shift of the PPS boundary toward the virtual body (Lenggenhager et al., 2007; Noel et al., 2015). Consciousness is characterized by a unity of experience in which information from multiple sensory modalities is integrated and bound together (Bayne, 2002; James et al., 1981) and, accordingly, current theories of consciousness postulate that integration of information, including unconscious stimuli, is critical for perceptual awareness (Baars, 2002; Mudrik et al., 2014; Tononi, 2008). Recent work has shown that consciously perceived stimuli can be integrated with subliminal stimuli (Alsius & Munhall, 2013; Lunghi et al., 2010; e.g. Lunghi et al., 2014; Salomon, Galli, et al., 2015; Salomon, Kaliuzhna, et al., 2015; Salomon, Lim, Herbelin, et al., 2013; Zhou et al., 2010). The present data show that unconscious multisensory integration also extends to a more complex and specific form of conscious content (Dehaene & Changeux, 2011; Faivre et al., 2015; Gallagher, 2000), i.e., self-consciousness targeted experimentally through BSC. Thus, we provide the first experimental support to the idea that the multisensory integrative processes underlying BSC are enabled in the absence of stimulus awareness. The present findings show that the phenomenological content of self-consciousness is based on unconscious integration of bodily multisensory signals. Thus, BSC is strongly grounded in the field of psychophysical consciousness studies, suggesting that even more comprehensive notions of self-consciousness may follow similar principles.

## 5. Author Contributions

R.S., J.P.N., A.S., and O.B., conceived of the experiments, which were performed by R.S., J.P.N., M.L., and analyzed by R.S., J.P.N., and A.S. N.F., and T.M. provided valuable analysis tools and conceptual contributions to the manuscript, which was written by R.S., and A.S. All authors edited and approved the final version of the manuscript.

## References

Alsius, A., & Munhall, K. G. (2013). Detection of audiovisual speech correspondences without visual awareness. Psychological Science, 24(4), 423–431.

Arzi, A., Shedlesky, L., Ben-Shaul, M., Nasser, K., Oksenberg, A., Hairston, I. S., & Sobel, N. (2012). Humans can learn new information during sleep. Nature Neuroscience, 15(10), 1460–1465.

Baars, B. J. (2002). The conscious access hypothesis: origins and recent evidence. Trends in Cognitive Sciences, 6(1), 47–52.

Bayne, T. (2002). The unity of consciousness.

Blanke, O. (2012). Multisensory brain mechanisms of bodily self-consciousness. Nat Rev Neurosci, 13(8), 556–571.

Blanke, O., Landis, T., Spinelli, L., & Seeck, M. (2004). Out-of-body experience and autoscopy of neurological origin. Brain, 127(2), 243.

Blanke, O., & Metzinger, T. (2009). Full-body illusions and minimal phenomenal selfhood. Trends in Cognitive Sciences, 13(1), 7–13.

Blanke, O., Ortigue, S., Landis, T., & Seeck, M. (2002). Stimulating illusory own-body perceptions. Nature, 419(6904), 269–270.

Blanke, O., Slater, M., & Serino, A. (2015). Behavioral, Neural, and Computational Principles of Bodily Self-Consciousness. Neuron, 88(1), 145–166.

Canzoneri, E., Magosso, E., & Serino, A. (2012). Dynamic sounds capture the boundaries of peripersonal space representation in humans. PLoS One, 7(9), e44306.

Damasio, A. (2012). Self comes to mind: constructing the conscious brain: Random House Digital, Inc.

Dehaene, S., & Changeux, J. P. (2011). Experimental and theoretical approaches to conscious processing. Neuron, 70(2), 200–227.

Dehaene, S., & Naccache, L. (2001). Towards a cognitive neuroscience of consciousness: basic evidence and a workspace framework. Cognition, 79(1), 1–37.

Ehrsson, H. H. (2007). The experimental induction of out-of-body experiences. Science, 317(5841), 1048–1048.

Ehrsson, H. H. (2012a). 43 The Concept of Body Ownership and Its Relation to Multisensory Integration.

Ehrsson, H. H. (2012b). The Concept of Body Ownership and Its Relation to Multisensory Integration.

Faivre, N., Mudrik, L., Schwartz, N., & Koch, C. (2014). Multisensory Integration in Complete Unawareness Evidence From Audiovisual Congruency Priming. Psychological Science, 25(11), 2006–2016.

Faivre, N., Salomon, R., & Blanke, O. (2015). Visual consciousness and bodily self-consciousness. Current opinion in neurology, 28(1), 23–28.

Gallagher, S. (2000). Philosophical conceptions of the self: Implications for cognitive science. Trends in Cognitive Sciences, 4(1), 14–21.

Graziano, M. S., & Cooke, D. F. (2006). Parieto-frontal interactions, personal space, and defensive behavior. Neuropsychologia, 44(6), 845–859.

Graziano, M. S., Hu, X. T., & Gross, C. G. (1997). Visuospatial properties of ventral premotor cortex. J Neurophysiol, 77(5), 2268–2292.

Graziano, M. S., Hu, X. T., & Gross, C. G. (1997). Visuospatial properties of ventral premotor cortex. Journal of neurophysiology, 77(5), 2268–2292.

Ionta, S., Heydrich, L., Lenggenhager, B., Mouthon, M., Fornari, E., Chapuis, D., Gassert, R., & Blanke, O. (2011). Multisensory Mechanisms in Temporo-Parietal Cortex Support Self-Location and First-Person Perspective. Neuron, 70(2), 363–374.

James, W., Burkhardt, F., Bowers, F., & Skrupskelis, I. (1981). The principles of psychology: Harvard Univ Pr.

Ladavas, E., & Serino, A. (2008). Action-dependent plasticity in peripersonal space representations. Cogn Neuropsychol, 25(7-8), 1099–1113.

Lenggenhager, B., Mouthon, M., & Blanke, O. (2009). Spatial aspects of bodily self-consciousness. Consciousness and Cognition, 18(1), 110–117.

Lenggenhager, B., Tadi, T., Metzinger, T., & Blanke, O. (2007). Video ergo sum: manipulating bodily self-consciousness. Science, 317(5841), 1096.

Lunghi, C., & Alais, D. (2013). Touch interacts with vision during binocular rivalry with a tight orientation tuning. PLoS ONE, 8(3), e58754.

Lunghi, C., Binda, P., & Morrone, M. C. (2010). Touch disambiguates rivalrous perception at early stages of visual analysis. Current Biology, 20(4), R143–R144.

Lunghi, C., Morrone, M. C., & Alais, D. (2014). Auditory and Tactile Signals Combine to Influence Vision during Binocular Rivalry. The Journal of Neuroscience, 34(3), 784–792.

Makin, T. R., Holmes, N. P., & Ehrsson, H. H. (2008). On the other hand: dummy hands and peripersonal space. Behavioural Brain Research, 191(1), 1–10.

Maruya, K., Yang, E., & Blake, R. (2007). Voluntary action influences visual competition. Psychological Science, 18(12), 1090–1098.

Meredith, M. A., & Stein, B. E. (1986). Visual, auditory, and somatosensory convergence on cells in superior colliculus results in multisensory integration. Journal of neurophysiology, 56(3), 640–662.

Metzinger, T. (2004). Being no one: The self-model theory of subjectivity: mit Press.

Miles, W. R. (1930). Ocular dominance in human adults. The Journal of General Psychology, 3(3), 412–430.

Mudrik, L., Faivre, N., & Koch, C. (2014). Information integration without awareness. Trends in Cognitive Sciences.

Noel, J.-P., Wallace, M., & Blake, R. (2015). Cognitive Neuroscience: Integration of Sight and Sound outside of Awareness? Current Biology, 25(4), R157–R159.

Noel, J. P., Grivaz, P., Marmaroli, P., Lissek, H., Blanke, O., & Serino, A. (2014). Full body action remapping of peripersonal space: The case of walking. Neuropsychologia.

Noel, J. P., Pfeiffer, C., Blanke, O., & Serino, A. (2015). Peripersonal space as the space of the bodily self. Cognition, 144, 49–57.

Petkova, V. I., & Ehrsson, H. H. (2008). If I were you: perceptual illusion of body swapping. PLoS ONE, 3(12), e3832.

Petkova, V. I., Khoshnevis, M., & Ehrsson, H. H. (2011). The perspective matters! Multisensory integration in ego-centric reference frames determines full-body ownership. Frontiers in psychology, 2.

Rizzolatti, G., Fadiga, L., Fogassi, L., & Gallese, V. (1997). NEUROSCIENCE: Enhanced: The Space Around Us 10.1126/science.277.5323.190. Science, 277(5323), 190–191.

Salomon, R., Galli, G., Łukowska, M., Faivre, N., Ruiz, J. B., & Blanke, O. (2015). An invisible touch: Body-related multisensory conflicts modulate visual consciousness. Neuropsychologia.

Salomon, R., Kaliuzhna, M., Herbelin, B., & Blanke, O. (2015). Balancing awareness: Vestibular signals modulate visual consciousness in the absence of awareness. Consciousness and Cognition, 36, 289–297.

Salomon, R., Lim, M., Herbelin, B., Hesselmann, G., & Blanke, O. (2013). Posing for awareness: Proprioception modulates access to visual consciousness in a continuous flash suppression task. Journal of Vision, 13(7).

Salomon, R., Lim, M., Pfeiffer, C., Gassert, R., & Blanke, O. (2013). Full body illusion is associated with widespread skin temperature reduction. Frontiers in behavioral neuroscience.

Serino, A., Canzoneri, E., Marzolla, M., di Pellegrino, G., & Magosso, E. (2015). Extending peripersonal space representation without tool-use: evidence from a combined behavioral-computational approach. Front Behav Neurosci, 9, 4.

Serino, A., Noel, J.P., Galli, G., Marmaroli P., Lissek, H., Blanke, O. (In press). Body parts-centered versus and full body-centered peripersonal space representations. Scientific Reports.

Stein, B. E., & Stanford, T. R. (2008). Multisensory integration: current issues from the perspective of the single neuron. Nat Rev Neurosci, 9(4), 255–266.

Teneggi, C., Canzoneri, E., di Pellegrino, G., & Serino, A. (2013). Social modulation of peripersonal space boundaries. Curr Biol, 23(5), 406–411.

Tononi, G. (2008). Consciousness as integrated information: a provisional manifesto. The Biological Bulletin, 215(3), 216–242.

Tsakiris, M. (2010). My body in the brain: a neurocognitive model of body-ownership. Neuropsychologia, 48(3), 703–712.

Tsuchiya, N., & Koch, C. (2005). Continuous flash suppression reduces negative afterimages. Nature Neuroscience, 8(8), 1096–1101.

Yang, E., & Blake, R. (2012). Deconstructing continuous flash suppression. Journal of Vision, 12(3), 8.

Zhou, W., Jiang, Y., He, S., & Chen, D. (2010). Olfaction Modulates Visual Perception in Binocular Rivalry. Current Biology, 20(15), 1356–1358.

